# Exome Sequencing in Individuals with Isolated Biliary Atresia

**DOI:** 10.1101/831768

**Authors:** Ramakrishnan Rajagopalan, Ellen A. Tsai, Christopher M. Grochowski, Susan M. Kelly, Kathleen M. Loomes, Nancy B. Spinner, Marcella Devoto

## Abstract

Biliary atresia (BA) is a severe pediatric liver disease resulting in necroinflammatory obliteration of the extrahepatic biliary tree. BA presents within the first few months of life as either an isolated finding or with additional syndromic features. The etiology of isolated BA is unknown, with evidence for infectious, environmental, and genetic risk factors described. However, to date, there are no definitive causal genes identified for isolated BA in humans, and the question of whether single gene defects play a major role remains open. We performed exome-sequencing in 100 North American patients of European descent with isolated BA (including 30 parent-child trios) and considered several experimental designs to identify potentially deleterious protein-altering variants that may be involved in the disease. In a case-only analysis, we did not identify genes with variants shared among more than two probands, and burden tests of rare variants using a case-case control design did not yield significant results. In the trio analysis of 30 simplex families (patient and parent trios), we identified 66 *de novo* variants in 66 genes including a nonsense variant, p.(Cys30Ter), in the gene *STIP1. STIP1* is a co-chaperone for the heat-shock protein, *HSP90AA1*, and has been shown to have diverse functions in yeast, flies and mammals, including stress-response.

**Conclusion:** Our results do not support the hypothesis that a simple genetic model is responsible for the majority of cases of isolated BA. Our finding of a *de novo* mutation in a candidate gene for BA (*STIP1*) linked to evolutionarily conserved stress responses suggests further exploration of how genetic susceptibility and environmental exposure interact to cause BA is warranted.

## Introduction

Biliary atresia (BA) is an obstructive cholangiopathy with initial symptoms arising during the first days to weeks of life. BA occurs as an isolated finding in 85% of affected individuals, and with additional syndromic features (heterotaxy and/or other congenital birth defects) in 15%^1^. The incidence of BA varies across different populations, with estimates ranging from 1 in 5,000 to 1 in 14,000 live births^2^. BA presents clinically with neonatal cholestasis, elevated bilirubin and liver enzymes, although the differential diagnosis suggested by these findings is extensive. The diagnosis of BA is suggested by features of biliary obstruction on liver histology, including bile duct proliferation, portal tract expansion and bile plugs. The diagnosis is confirmed upon intraoperative cholangiogram showing lack of patency of the extrahepatic biliary tree.

Although the etiology of BA is not clear, a genetic component is supported by multiple lines of evidence. These include the presence of familial cases^3–5^, case reports of individuals with syndromic BA with likely causal genes identified (including *FOXA2^6^, CFC1*^7,8^, *ZEB2*^9^, *ZIC3*^10^, *HNF1B*^11^ and *PKD1L1*^12^), animal models of BA involving the genes *Sox17*^13^, *Foxm1*^14^, *Invs*^15^, *Onecutl*^16^, and candidate susceptibility genes including *ADD3*^17^, *GPC1*^18^, *EFEMP1*^19^ and *ARF6*^20^ identified by genome-wide association studies (GWAS). Genes implicated in syndromic BA function in bile-duct development and the establishment of left-right symmetry. A follow-up study of *GPC1* using a zebrafish model confirmed that suppressing *GPC1* expression gives rise to biliary defects^21^. A GWAS identified SNPs associated with BA upstream of *XPNPEP1* and *ADD3* in a Chinese cohort of isolated BA patients^17^, and this association has been confirmed in both a Thai^22^ and an European-American cohort^23^. In a study of 499 isolated and syndromic BA patients and 1928 controls, our group identified a GWAS signal within 2p16.1 implicating common variants in the gene *EFEMP1*^19^. Another genome-wide association study of 63 BA patients and 1907 controls identified two SNPs downstream of the gene *ARF6* associated with isolated BA^20^, although we could not replicate this finding in our study. However, to date, no genes have been definitively identified as a cause of isolated BA. Environmental exposure to virus or toxins has long been proposed as a hypothesis for perinatal form of BA^2^. But the genetic susceptibility factors linking such exposure and BA are yet to be identified.

In this work, we hypothesized that rare and novel, deleterious, protein-altering variants transmitted in a Mendelian fashion are involved in the etiology of isolated BA, and patients with isolated BA are likely to be enriched with such variants. To test this hypothesis, we considered three experimental designs (*Table 1*). First, we conducted a case-only analysis where we looked for potentially deleterious novel variants. In the second design, we conducted a case-control analysis comparing the cumulative frequency of rare variants in the two groups. Finally, we performed trio analysis on a subset of 30 probands to identify genes with rare, protein-altering *de novo* and compound heterozygous or homozygous variants.

**Table 1:**
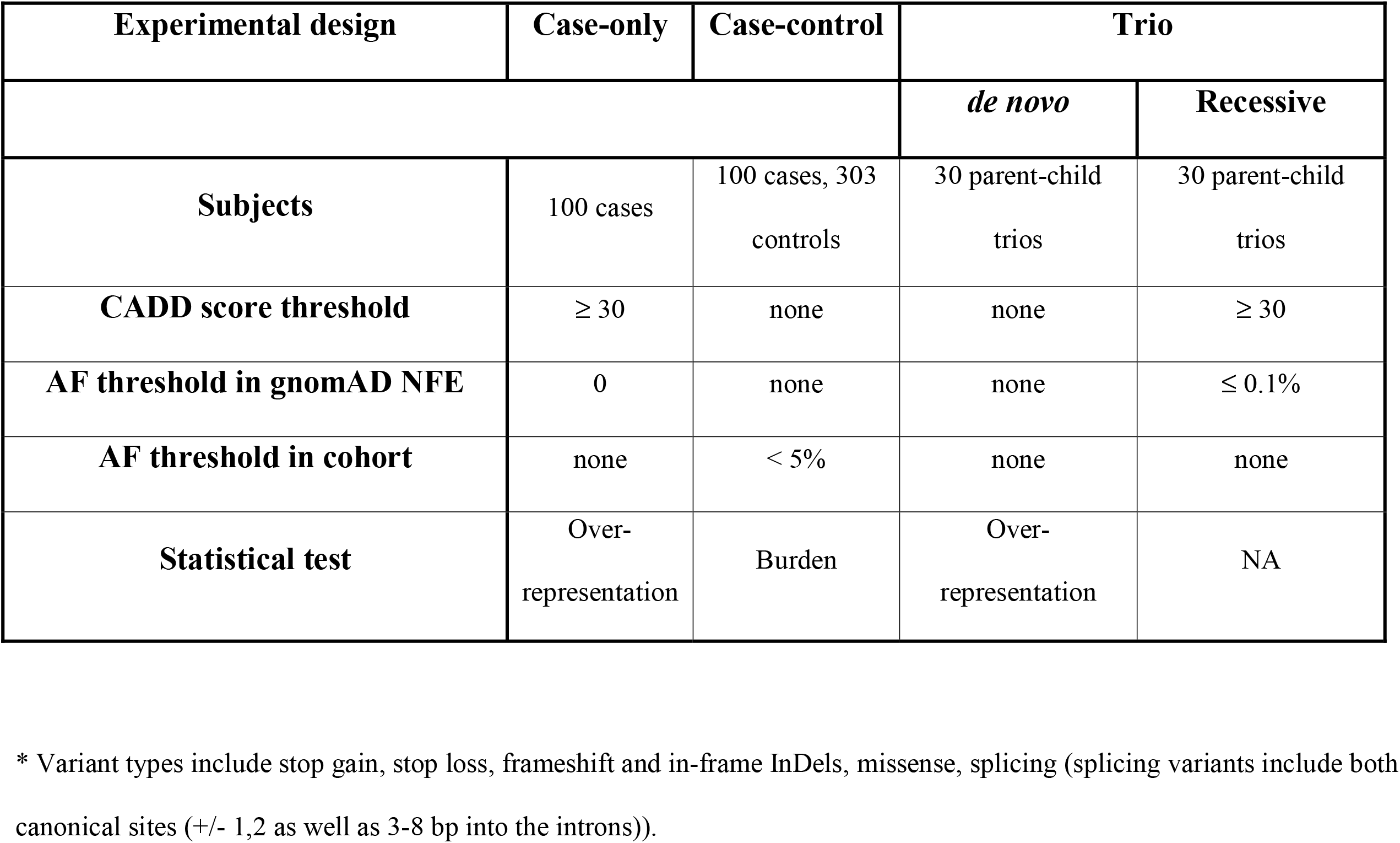
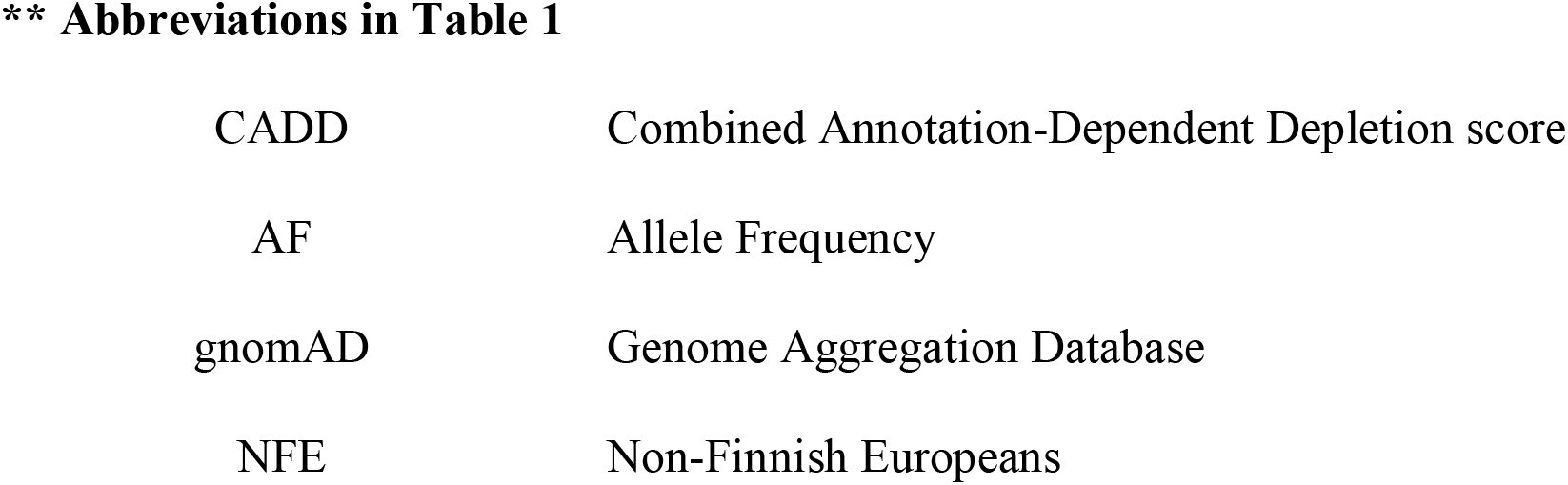
Experimental designs used in this study

| Experimental design | Case-only | Case-control | Trio |  |
| --- | --- | --- | --- | --- |
|  |  |  | *de novo* | Recessive |
| **Subjects** | 100 cases | 100 cases, 303 controls | 30 parent-child trios | 30 parent-child trios |
| **CADD score threshold** | $\geq 30$ | none | none | $\geq 30$ |
| **AF threshold in gnomAD NFE** | 0 | none | none | $\leq 0.1\%$ |
| **AF threshold in cohort** | none | $< 5\%$ | none | none |
| **Statistical test** | Over-representation | Burden | Over-representation | NA |

\* Variant types include stop gain, stop loss, frameshift and in-frame InDels, missense, splicing (splicing variants include both canonical sites (+/- 1,2 as well as 3-8 bp into the introns)).

**\*\* Abbreviations in Table 1**
|  |  |
| --- | --- |
| CADD | Combined Annotation-Dependent Depletion score |
| AF | Allele Frequency |
| gnomAD | Genome Aggregation Database |
| NFE | Non-Finnish Europeans |

## Patients and Methods

### Patients with isolated BA and controls

We studied a cohort of 100 self-reported white, non-Hispanic patients with isolated BA. The 100 patients were chosen from a larger cohort of 1063 BA cases based on their self-reported race and ethnicity and availability of parental DNA samples. Ninety-nine participants were enrolled in the NIDDK-funded Childhood Liver Disease Research Network (ChiLDReN) at one of the sixteen participating institutions under Institutional Review Board (IRB) approved protocols (IRB 04-003655 and 06-004814). A set of dizygotic twins were consented into the approved IRB study at the Children’s Memorial Hospital, Chicago, IL and only one of them were part of the cohort as the scope of the current work was to analyze unrelated individuals. Informed consent in wirting was obtained from each patient enrolled in this study. The diagnosis of BA was made by clinical presentation, liver histology, and an intraoperative cholangiogram. Most patients also had BA confirmed by examination of the biliary remnant from a Roux-en-Y hepatic portoenterostomy (also known as a Kasai operation). DNA samples were extracted from whole blood or lymphoblastoid cell lines. For a subset of patients (n = 30), DNA samples were available from both parents.

Controls were individuals recruited from two different studies at the Children’s Hospital of Philadelphia, PediSeq (the CHOP cohort of the Clinical Sequencing and Exploratory Research (CSER) consortium^24^, and VEO-IBD (Very-Early Onset Inflammatory Bowel Disease)^25^, and enrolled in institution-approved IRB protocols (IRB 12-009169; 14-010826). Controls from the PediSeq dataset (n=158) fall into four different disease cohorts (cardiovascular, hearing loss, mitochondrial and intellectual disability) but have no known liver disease. Controls from the VEO-IBD cohort (n=145) were the unaffected parents of individuals with VEO-IBD. The patient recruitment, methods, and experimental protocols used in these studies were performed in accordance with relevant guidelines and regulations set by the aforementioned institutions. This study was approved by the ethics committee at the Children’s Hospital of Philadelphia.

### Exome sequencing (ES)

Genomic DNA (3-5 ug) was supplied to Perkin Elmer, Inc. (Perkin-Elmer, Branford, CT) for whole exome sequencing of the BA patients and their parents. The DNA was amplified and enriched using the Agilent SureSelect V4+UTR 71Mb All Exon Capture Kit (Agilent Technologies, Santa Clara, CA). Sequencing was performed on the Illumina HiSeq 2000 (Illumina, San Diego, CA) at the Perkin-Elmer sequencing service center. Control samples were sequenced using Agilent SureSelect V4 51Mb Exon Capture Kit (Agilent Technologies, Santa Clara, CA) on the Illumina HiSeq 2000 (Illumina, San Diego, CA) at the internal sequencing core at CHOP (BGI@CHOP, Philadelphia, PA). All of the exome sequencing data was produced with paired-end reads of 100 base-pairs in length (2 x 100bp). The unaligned FASTQ reads of all samples were returned to us for read analysis and downstream processing as detailed below.

### Variant calling and annotation

Raw sequencing reads were mapped to the GRCh37.69 reference genome using the Burrows-Wheeler Aligner^26^ to produce an aligned BAM file. BAM files were realigned around known indels using IndelRealigner in the Genome Analysis ToolKit software (GATK) and base quality scores were recalibrated using the BaseQualityScoreRecalibrator in GATK^27^ Variant calling was performed using HaplotypeCaller and jointly genotyped using the GenotypeGVCFs command in GATK. The initial variant quality filtering was done using the VariantQualityScoreRecalibrator (VQSR) in GATK. SNPEff^28^ was used to annotate the variants with their expected functional consequence. Other annotations such as allele frequencies in non-Finnish Europeans (NFE) from the gnomAD variant database, and deleteriousness scores such as CADD (Combined Annotation Dependent Depletion version 1.4)^29^ were added using the Gemini software^30^. We manually reviewed all variants found in the *de novo* analysis, and variant clusters with overlapping genomic positions that could be indicative of technical artifacts^31^.

### Population stratification, relatedness and sex check

The Peddy^32^ software tool was used to infer genetic ancestry from the exome sequencing data by comparing our samples against the samples from the 1000 Genomes Project Phase 3^33^. All samples from BA patients and their parents clustered with the European populations in the 1000 Genomes Project. We also confirmed the sex of the participants based on heterozygosity of X chromosome variants. Pairwise relatedness (estimated by the kinship coefficient) between samples was investigated using the KING^34^ software and confirmed that there were no related individuals in the entire cohort beside the known parent-child pairs used in the trio analysis.

### Variant prioritization

We filtered the variants identified from the ES for stop gain, stop loss, frameshift and inframe insertions/deletions (InDels), missense and splicing variants. We excluded variants overlapping with segmental duplications and variants with call rate less than 90% from all analyses. In the case-only analysis, we included novel and potentially deleterious variants, defined as variants that were not present in the Non-Finnish European (NFE) population of the gnomAD genomes and exomes and had a phred-scaled CADD score of at least 30, representing the top 0.1% of the potentially deterious variants. For the burden tests in the case-control experimental design, all protein-altering variants with a cohort (cases and controls combined) minor allele frequency (MAF) of <5% were included. We did not apply a CADD score threshold for this analysis. For the *de novo* analysis, we considered all variants irrespective of their frequency or the CADD score. For the analysis under the recessive model, we retained variants with an allele frequency in gnomAD NFE less than or equal to 0.1% and a CADD score of at least 30. Details pertaining to variant prioritization are provided in *Table 1*.

### Autosomal copy-number variant (CNV) analysis

Mean depth of coverage for each individual exonic interval in the autosomes was computed using the DepthOfCoverage routine in the GATK package and copy number variants were called using XHMM following the protocol described elsewhere^35^. We excluded CNVs with quality score (Q_SOME) less than 60 to retain only high quality CNVs. To identify novel CNVs, we excluded any variant found in the Database of Genomic Variants (DGV)^36^ and the ExAC^37^ We did not look for CNVs in the sex chromosomes as the ExAC dataset only had calls from autosomes.

### Gene, pathway and gene-set definitions

For all the analyses, we used refFlat_hg19 gene definitions from the UCSC genome browser^38^. We used pathway definitions including KEGG (n = 293), Panther (n = 112), Reactome (n = 1530), WikiPathways (n = 437) and BioCarta (n = 237), and annotation gene-sets from Gene Ontology (GO) Biological Process (n = 5192), Molecular Functions (n = 1136) and Cellular Components (n=641) downloaded from the Enrichr website.^39^

### Case-only and trio analyses: Over representation tests

To determine if our lists of candidate genes from the case-only and trio analyses were enriched for specific gene-sets or pathways, we performed over-representation analysis using the R package enrichR^39^. This software essentially counts the number of genes with a given annotation that are present in our lists of candidate genes and genome-wide, and compares it to the number of genes that are not associated with the same annotation in our lists and genome-wide, using a Fisher exact test. Over representation tests were carried out on gene-sets and pathways.

### Case-control analysis: Rare variant burden tests

To test if a gene, gene-set or pathway contained significantly more cases with at least one rare variant (MAF < 5%) compared to controls, we performed burden tests using the Combined and Multivariate Collapsing (CMC) approach^40^ implemented in the exactCMC procedure from the software package rvtests^41^. Burden tests were carried out at different levels including 1) the gene level; 2) the gene-set level (e.g. collection of genes with similar function or structure or cellular localization, such as sodium channel genes or membrane bound receptors); and 3) the pathway level, which includes genes that function in a specific biological pathway (e.g. WNT or NOTCH signaling). The number of cases and controls with and without at least one rare variant in a given gene, gene-set, or pathway were compared using a Fisher exact test.

### Trio analysis

Trio analysis was performed using Gemini software^30^ in a subset of 30 probands for whom DNA from both parents was available for ES. *De novo* variants were identified using the command *‘de_novo’* which essentially identifies SNVs and indels occuring in proband only and not in the parents. Further, we filtered for exonic, protein-altering and splice-site variants. Compound heterozygous variants were identified using the command ‘*comp_het*’ which identifies two heterozygous variants in the same gene inherited from different parents. Homozygous variants were identified using the ‘*autosomal_recessive*’ command which identifies rare variants transmitted to the proband from both parents.

To test if the total number of *de novo* protein-altering (missense and loss-of-function) variants identified in our cohort was significantly larger than expected, we used the R-package *denovolyzeR*^42^. The *denovolyzeByClass* function implemented in the denovolyzeR package computes the significance of the observed number of exonic variants compared to the pre-computed *de novo* mutation rates for all classes of exonic variants.

### Statistical significance

For gene level burden tests, the raw p-values were corrected for multiple testing using a Bonferroni correction for the total number of genes tested (n=16,393). For the pathway based analyses, the raw P-values were corrected for multiple testing using a Bonferroni correction for the total number of pathway definitions (n=2609). For the gene-set based analyses, we corrected for the total number of gene-sets used in the analysis (n = 6969). We considered a test to be statistically significant if the Bonferroni-adjusted p-value was less than or equal to 0.05.

## Results

### Case-only analysis

In the case-only analysis we identified a total of 496,654 potentially deleterious novel variants in the 100 probands. These included variants that passed the quality filters, of which 63,473 single nucleotide variants (SNV) or small InDels were found to be protein-altering changes (stop gain, stop loss, frameshift and in-frame InDels, missense and splicing variants). Of these, 332 variants in 326 genes were novel (not in the NFE population in gnomAD exomes and genomes) and had a phred-scaled CADD score of 30 or more. This formed our variant dataset for the case-only analyses. Almost all of the variants (331/332) were present in heterozygosity in a single individual, and 96 out of the 100 individuals in the cohort had at least one potentially deleterious novel variant. Six genes had two different variants in two different individuals (*MAST2, WDR35, DNAH5, FBN2, SCUBE2 and PITPNM3*). Over-representation analysis did not yield any significant gene, pathway or gene-set after correcting for multiple testing. Copy-number variant analysis did not reveal any novel variant in the probands after filtering for variants present in the population databases ExAC and DGV. Details of the SNVs and InDels with annotations are provided in the *Supp. Table 1*.

### Case-control analysis

In the second experimental design, we conducted burden tests for rare variants (MAF < 5%) by comparing their cumulative number in a gene, gene-set or pathway in the 100 cases against the 303 controls. After correcting for the multiple testing, we did not identify any gene, gene-set or a pathway with a statistically significant excess of rare variants in the cases compared to the controls.

### Trio analysis

In the final experimental design, we sequenced the parents of a subset of the patients (n = 30) to identify *de novo* variants in the probands under the hypothesis that they are more likely to be involved in the etiology of the disease. We identified a total of 66 *de novo,* protein-altering and splice variants in 66 different genes in 25 out of 30 individuals, with a mean of 2.2 variants per individual (range: 0-7; median 3). We did not identify any *de novo* variants in five individuals. All the *de novo* variants were manually verified in the BAM file and a small subset of them were validated using Sanger sequencing (n = 4). There were 58 missense, 3 frameshift indels, 3 nonsense, and 2 splice-site variants. A prioritized list of 14 *de novo* variants are provided in *Table 2* and the list of all *de novo* variants are provided in the *Supp. Table 2*. The number of protein-altering *de novo* variants identified in this cohort was significantly larger than the expected number of such variants in a cohort of 30 individuals (expected = 21.5; observed = 66; p-value = 1.13×10^−14^). This signal was mainly driven by the missense variants (expected = 18.9; observed = 58; p-value = 4.15e^−13^), while the loss-of-function variants had a modest significance (expected = 2.6; observed = 8; p-value = 5.61×10^−3^). Of the eight putative *de novo* loss-of-function variants, a premature stop-gain in the gene *STIP1* and a spliceacceptor variant in the gene *REV1* were the most interesting as they are known to be depleted for and intolerant to loss-of-function in the general population with a pLI score of 1. The pLI or probability of loss-of-function intolerance is a gene-level measure estimated by comparing the expected number of loss-of-function variants against the observed in a population. The pLI score ranges from 0 to 1 with the highest value being the most intolerant. Over-representation analysis of the 66 genes did not yield significant results after correcting for multiple testing. We did not identify any compound heterozygous or homozygous variants at a maximum alternate allele frequency threshold of 0.1% and a phred-scaled CADD score of 30.

**Table 2:**
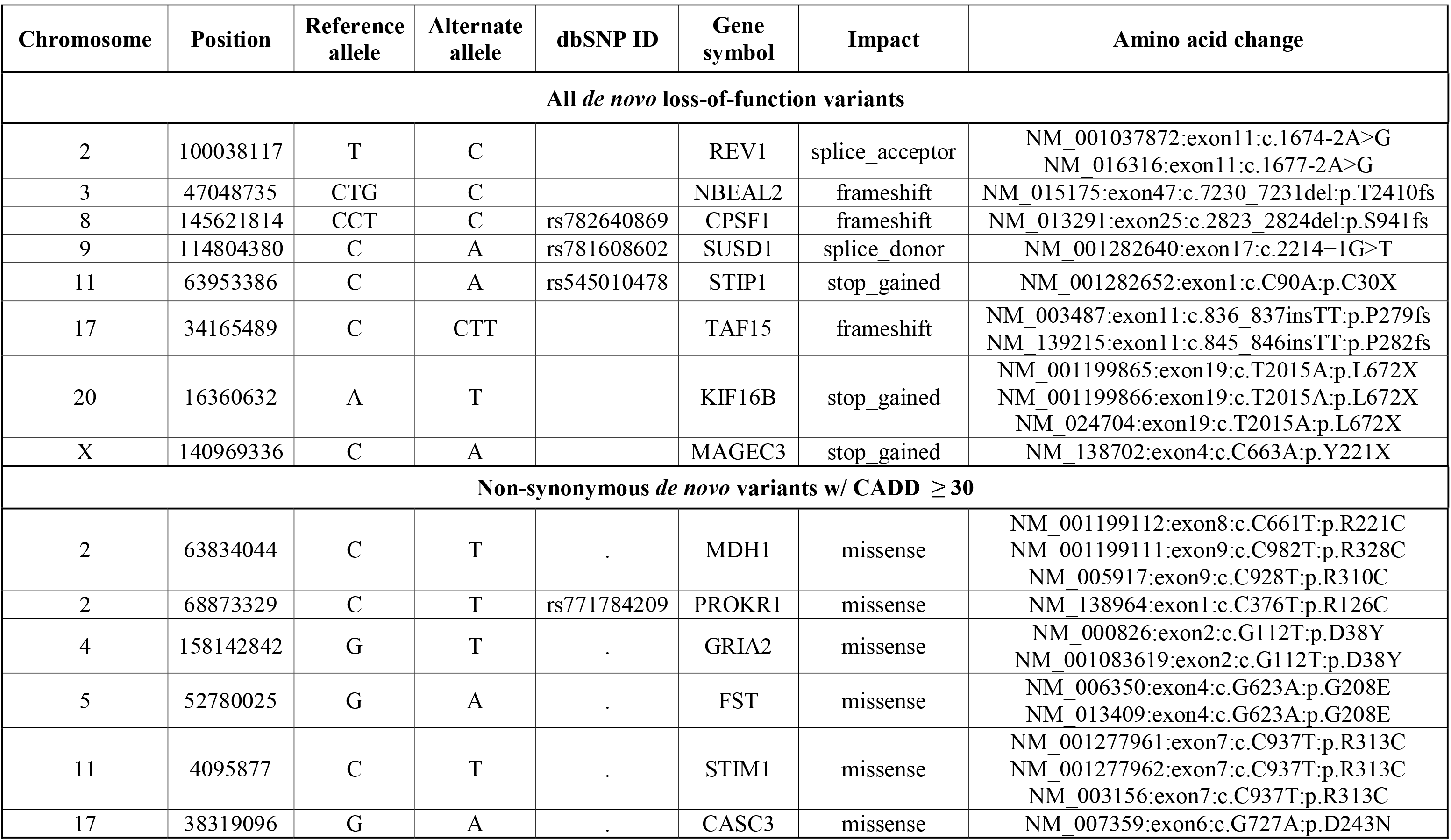
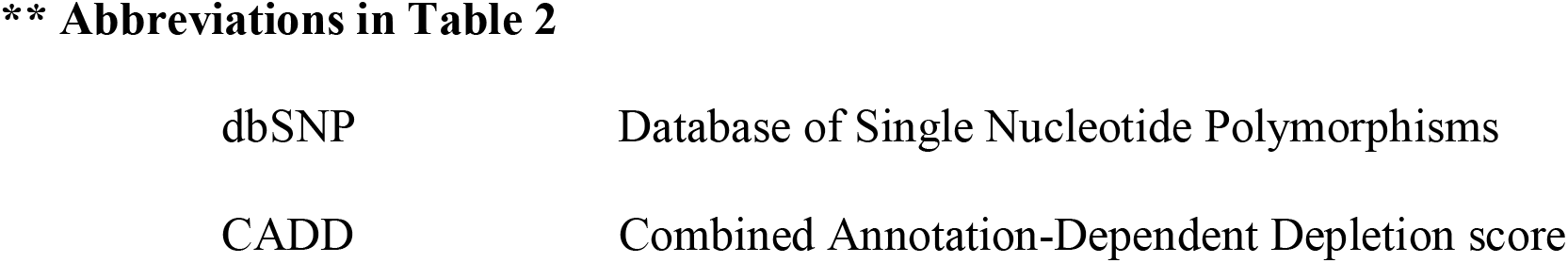
Prioritized list of *de novo* variants identified in the trio analysis of 30 parentchild trios. This table lists all the loss-of-function, splice variants and missense variants with a CADD score greater than 30.

| Chromosome | Position | Reference allele | Alternate allele | dbSNP ID | Gene symbol | Impact | Amino acid change |
| --- | --- | --- | --- | --- | --- | --- | --- |
| **All *de novo* loss-of-function variants** |  |  |  |  |  |  |  |
| 2 | 100038117 | T | C |  | REV1 | splice_acceptor | NM_001037872:exon11:c.1674-2A>G<br>NM_016316:exon11:c.1677-2A>G |
| 3 | 47048735 | CTG | C |  | NBEAL2 | frameshift | NM_015175:exon47:c.7230_7231del:p.T2410fs |
| 8 | 145621814 | CCT | C | rs782640869 | CPSF1 | frameshift | NM_013291:exon25:c.2823_2824del:p.S941fs |
| 9 | 114804380 | C | A | rs781608602 | SUSD1 | splice_donor | NM_001282640:exon17:c.2214+1G>T |
| 11 | 63953386 | C | A | rs545010478 | STIP1 | stop_gained | NM_001282652:exon1:c.C90A:p.C30X |
| 17 | 34165489 | C | CTT |  | TAF15 | frameshift | NM_003487:exon11:c.836_837insTT:p.P279fs<br>NM_139215:exon11:c.845_846insTT:p.P282fs |
| 20 | 16360632 | A | T |  | KIF16B | stop_gained | NM_001199865:exon19:c.T2015A:p.L672X<br>NM_001199866:exon19:c.T2015A:p.L672X<br>NM_024704:exon19:c.T2015A:p.L672X |
| X | 140969336 | C | A |  | MAGEC3 | stop_gained | NM_138702:exon4:c.C663A:p.Y221X |
| **Non-synonymous *de novo* variants w/ CADD ≥ 30** |  |  |  |  |  |  |  |
| 2 | 63834044 | C | T | . | MDH1 | missense | NM_001199112:exon8:c.C661T:p.R221C<br>NM_001199111:exon9:c.C982T:p.R328C<br>NM_005917:exon9:c.C928T:p.R310C |
| 2 | 68873329 | C | T | rs771784209 | PROKR1 | missense | NM_138964:exon1:c.C376T:p.R126C |
| 4 | 158142842 | G | T | . | GRIA2 | missense | NM_000826:exon2:c.G112T:p.D38Y<br>NM_001083619:exon2:c.G112T:p.D38Y |
| 5 | 52780025 | G | A | . | FST | missense | NM_006350:exon4:c.G623A:p.G208E<br>NM_013409:exon4:c.G623A:p.G208E |
| 11 | 4095877 | C | T | . | STIM1 | missense | NM_001277961:exon7:c.C937T:p.R313C<br>NM_001277962:exon7:c.C937T:p.R313C<br>NM_003156:exon7:c.C937T:p.R313C |
| 17 | 38319096 | G | A | . | CASC3 | missense | NM_007359:exon6:c.G727A:p.D243N |

**\*\* Abbreviations in Table 2**
|  |  |
| --- | --- |
| dbSNP | Database of Single Nucleotide Polymorphisms |
| CADD | Combined Annotation-Dependent Depletion score |

### Discussion

In this study, exome sequencing was used to identify genomic changes in a cohort of patients with isolated biliary atresia, and in spite of the utilization of several distinct strategies, we were unable to demonstrate an associated genetic change. We did identify a candidate gene by analysis of 30 trios, finding a *de novo* likely pathogenic variant in *STIP1*, which we hypothesize may result in susceptibility to toxin induced biliary disease.

BA presents as an isolated finding in 85% of cases, and in a syndromic form with laterality and/or other congenital malformations in 15%. Mouse models of BA (*Sox17, Onecut1*) and gene mutations in patients with syndromic forms of BA (*CFC1, ZIC3, FOXA2, PKD1L1*) suggest an etiology involving genetic factors that influence left-right symmetry, and bile duct development and morphology. Genome-wide association studies in patients with isolated BA have suggested candidate genes and genomic regions (*ADD3, XPNaaPEP1* in 10q25.1, *EFEMP1* in 2p16.1, *GPC1* in 2q37.3, *ARF6* in 14q21.3) but consistent proof of the involvement of each of these regions has been elusive. Biliary atresia as a consequence of exposure to environmental toxins is also proposed as a model for the perinatal form of the disease based on research into the natural occurance of BA in Australian sheep, which led to the discovery of an isoflavonoid (Biliatresone) found in specific plant species, later found to cause biliary defects in zebrafish^43^. RNA-seq experiments on Biliatresone treated and control fish identified several differentially expressed stress signaling pathways including the Glutathione antioxidant pathway, cellular redox response, and heat shock response^44^ However, the genetic susceptibility factors that might link exposure to these agents and biliary atresia are yet to be identified. In summary, from previous literature, there are genes that are known to cause syndromic forms of BA in humans and mice, genes that are associated with isolated BA identified via GWAS, and biological pathways that are perturbed in zebrafish treated with the toxin Biliatresone, which has been associated with biliary atresia in sheep. However, in spite of these clues, there has not been a causal gene identified for isolated BA in humans.

In this work, we performed exome sequencing in 100 North American unrelated patients of European ancestry with isolated BA and looked for novel or rare protein-altering variants that may explain the BA phenotype (strategies presented in *Table 1*). Such study designs have been successful in finding novel genes responsible for rare Mendelian diseases^45^. Fewer exome sequencing studies with sufficiently large sample sizes have been successfully employed in identifying associations with complex phenotypes^46,47^ This study was primarily designed with the goal of finding genes with rare and highly deleterious variants that may explain the occurrence of isolated BA. We considered three experimental designs including a case-only analysis to look for novel, potentially deleterious and protein-altering variants, a case-control analysis to look for an excessive burden of rare variants in the cases compared to the controls, and a trio analysis, to look for *de novo* and homozygous or compound heterozygous variants.

In the case-only design, we identified 332 novel and potentially deleterious protein-altering variants in our cohort of 100 individuals with isolated BA. The majority of the variants were found in a single individual in a heterozygous state. Six genes had variants in two individuals, and the variants were different from each other. This suggests that BA is unlikely to result from a single genetic change, although we cannot rule out extreme locus heterogeneity, with a small percentage of cases having a genetic etiology, perhaps in combination with an environmental insult. Case-control analysis did not yield any statistically significant result and this may suggest that there are no variants or genes with large enough effect sizes present in the cases as compared to the controls or simply the sample size is too small to identify such a difference between cases and controls.

Trio-based exome sequence analysis identified 66 *de novo* protein-altering variants in 66 genes in 25 out of 30 patients, a number significantly higher than expected (p-value = 4.15×10^−13^). All putative loss-of-function *de novo* variants and a prioritized list of *de novo* missense variants with a CADD score greater than or equal to 30 are provided in *Table 2*. Of all the genes with putative loss-of-function *de novo* variants, a splice-acceptor variant in the gene *REV1* and a premature stop in one of the transcripts in the gene *STIP1* were the most interesting for several reasons. These genes seem to be intolerant for loss-of-function variants with a perfect pLI score of 1. *STIP1* is a chaperone that assists in the transfer of proteins from *HSP70* to *HSP90,* which are an integral part of the heat shock response pathway and as mentioned below, *HSP90* down-regulation has been implicated in BA livers. Mass spectrometry experiments to identify differentially expressed proteins in 20 BA liver biopsies compared to 12 non-BA, neonatal cholestasis livers found that heat shock protein 90 (*HSP90*) was significantly down-regulated in BA livers, and was the most significantly altered protein. This suggests that *HSP90* could serve as a potential biomarker for BA^48^. We therefore hypothesized that a gene-environment interaction may explain isolated BA where a predisposing genetic factor exists and an external insult triggers the initiation of the disease. Based on the *de novo* variant found in the human exome data, the identification of the heatshock response pathway as differentially expressed in zebrafish, our collaborators performed a CRISPR/Cas9 based experiment to introduce a frameshift mutation in exon 1 of the zebrafish *stip1* gene to create heterozygous mutant fish. When treated with toxin, *stip1* heterozygous fish were highly sensitive to a low dose of biliatresone compared to the wild-type^49^. *REV1* encodes a protein similar to the *S. cerevisiae* mutagenesis protein *Rev1* and is known to be involved in DNA repair. REV1 is shown to be regulated by the heat-shock protein HSP90 in the DNA repair pathway^50^ and further functional studies are warranted to understand its role in BA.

Our study has several limitations including sample size (which is a function of the disease frequency) and the failure of exome sequencing to identify all possible disease causing genomic changes. The sample size was relatively small for statistical analyses such as rare variant burden tests and for the trio analysis whose aim was to identify genes with recurring *de novo* variants. Exome sequencing is a targeted capture experiment which focuses only on the exonic regions, and coverage might not be uniform through the coding parts of the genome. This limits our ability to study intronic regions and/or poorly targeted parts of the genome. Copy-number detection from exome sequencing is challenging and XHMM software tool used in this work is tuned for specificity rather than sensitivity. Finally, exome sequencing does not allow us to look for structural variation in these samples.

Together, our analysis of rare and novel coding variants does not support a simple genetic model in which a small number of genes are responsible for the majority of cases with isolated BA. However, the identification of a loss-of-function *de novo* variant in the gene *STIP1* and the existing evidence for its sensitivity to the biliatresone toxin in zebrafish opened new opportunities for further investigating BA as a consequence of environmental exposure. Potentially, other genes in the heat shock response pathway or other pathways identified in the zebrafish experiments could be considered as candidates.

## Acknowledgments

We would like to thank all the patients and their families who participated in this study; the ChiLDReN investigators, research coordinators and the Data Coordinating Center for the ChiLDReN network. We would like to acknowledge Dr. Judith Kelsen and Dr. Ian Krantz for providing exome sequencing data for the control samples. We also would like to acknowledge Dr. David Piccoli and Dr. Barbara Haber for their support. We would like to acknowledge Dr. Michael Pack and Dr. Xiao Zhao at the University of Pennsylvania for the collaborative work on zebrafish models.

## Author Contribudions

MD, KL, and NBS designed the experiments. RR and ET performed the computational and bioinformatics analysis. CG performed validation studies. KL and SK recruited patients. RR, ET, KL, NBS and MD were involved in the interpretation of the results. RR wrote the manuscript and all authors contributed in editing the manuscript.

## List of Tables

Table 1: Experimental designs and variant prioritization schemes used in this study

Table 2: Prioritized list of de novo variants identified in the trio analysis of 30 parent-child trios. This table lists all the loss-of-function, splice variants and missense variants with a CADD score greater than 30.

## Supporting Information

Supplementary Table 1: List of all the *de novo* variants identified

Supplementary Table 2: List of all the novel variants identified in the 100 probands

